# The infralimbic and prelimbic cortical areas bidirectionally regulate safety learning during normal and stress conditions

**DOI:** 10.1101/2023.05.05.539516

**Authors:** Ada C. Felix-Ortiz, Jaelyn M. Terrell, Carolina Gonzalez, Hope D. Msengi, Angelica R. Ramos, Miranda B. Boggan, Savannah M. Lopez-Pesina, Gabrielle Magalhães, Anthony Burgos-Robles

## Abstract

Safety learning is a critical function for behavioral adaptation, environmental fitness, and mental health. Animal models have implicated the prelimbic (PL) and infralimbic (IL) subregions of the medial prefrontal cortex (mPFC) in safety learning. However, whether these regions differentially contribute to safety learning and how their contributions become affected by stress still remain poorly understood. In this study, we evaluated these issues using a novel semi-naturalistic mouse model for threat and safety learning. As mice navigated within a test arena, they learned that specific zones were associated with either noxious cold temperatures (“threat”) or pleasant warm temperatures (“safety”). Optogenetic-mediated inhibition revealed critical roles for the IL and PL regions for selectively controlling safety learning during these naturalistic conditions. This form of safety learning was also highly susceptible to stress pre-exposure, and while IL inhibition mimicked the deficits produced by stress, PL inhibition fully rescued safety learning in stress-exposed mice. Collectively, these findings indicate that IL and PL bidirectionally regulate safety learning during naturalistic situations, with the IL region promoting this function and the PL region suppressing it, especially after stress. A model of balanced IL and PL activity is proposed as a fundamental mechanism for controlling safety learning.

**Visual Abstract:** 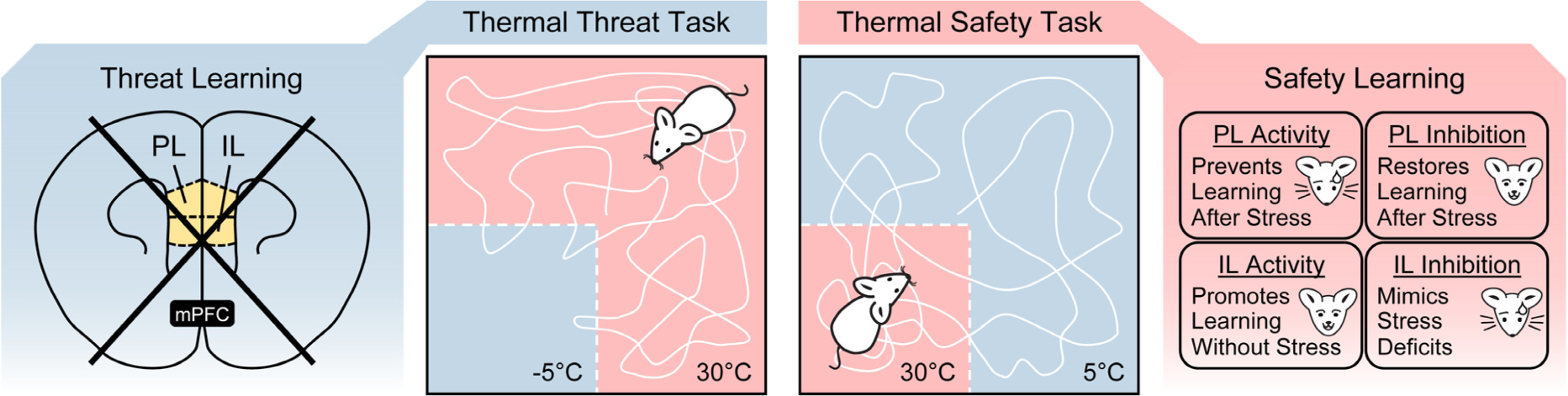

## Introduction

Safety learning allows flexible adaptation of behavior during threatening situations. While this function is critical for environmental fitness, it is also important for emotional regulation and mental health. For instance, safety learning is dysregulated in stress-related disorders, leading to excessive fear and anxiety, even in the absence of threat (Dunsmoor and Paz, 2015; Jovanovic et al., 2012). Yet, the neural mechanisms that normally control safety learning and how they get affected by stress to produce issues with emotional regulation still remain elusive.

Growing evidence indicates that safety learning requires similar neural networks to threat learning. Most models position the amygdala at the front and center to dissociate the environmental cues that predict threat versus safety (Genud-Gabai et al., 2013; LeDoux and Daw, 2018; Sangha et al., 2013; Stujenske et al., 2014; Tovote et al., 2015; Wen et al., 2022). The hippocampus is also critical to distinguish contextual signals that predict threat versus safety (Meyer et al., 2019; Moscarello and Maren, 2018; Rozeske et al., 2015). Current models also highlight contributions of mPFC regions that can integrate amygdala and hippocampal inputs to promote higher-order functions such as decision making, conflict resolution, and memory updating for flexible adaptation of behavior during situations involving threat (Bravo-Rivera and Sotres-Bayon, 2020; Burgos-Robles et al., 2017; Corches et al., 2019; Fernandez-Leon et al., 2021; Sharpe and Killcross, 2014; Tashjian et al., 2021).

The mPFC in rodents is often divided into the PL and IL regions, which have been identified as key elements in the networks controlling distinct forms of threat and safety learning. Despite their anatomical proximity and similarities in cell composition (Anastasiades and Carter, 2021), PL and IL exhibit remarkable differences in function for controlling threat-related behavior (Sierra-Mercado et al., 2011; Vidal-Gonzalez et al., 2006). For instance, PL exhibits robust signals that promote the acquisition and expression of conditioned responses during threat-predicting cues (Burgos-Robles et al., 2009; Diehl et al., 2018; Gilmartin and McEchron, 2005; Laviolette et al., 2005). In contrast, IL exhibits strong signals that correlate with the extinction and inhibition of conditioned responses when threatening cues no longer predict danger (Burgos-Robles et al., 2007; Milad and Quirk, 2002; Russo and Parsons, 2022; Tao et al., 2021). Similar dissociations between PL and IL function have been reported during paradigms involving discrete cues that differentially predict threat and safety (Likhtik et al., 2014; Meyer and Bucci, 2014; Sangha et al., 2014). Thus, current models strongly argue that PL and IL contribute in opposite manners during situations involving threat to allow threat learning, safety learning, and behavioral flexibility (Odriozola and Gee, 2021; Sangha et al., 2020; Sotres-Bayon and Quirk, 2010). Yet, additional insights are needed to better understand the contributions of these regions during safety learning.

While the number of studies implicating the mPFC in dissociations between threat and safety continues to grow, the majority of previous studies have used cue discrimination tasks in which auditory cues predict either the delivery or omission of electric shock punishment. This raises the question of whether the mPFC is still relevant during more naturalistic situations. Interestingly, recent imaging studies in humans reported that mPFC regions signal thermal stimuli, presumably to evaluate their hedonic value (i.e., determine whether they are noxious or pleasant; Aizawa et al., 2019; Elman et al., 2018). Since thermal stimuli elicit defensive behavioral mechanisms in many animal species including humans, (Bokiniec et al., 2018; Kanai et al., 2022; Ofstad et al., 2011; Wang et al., 2021; Wessnitzer et al., 2008), thermal stimuli seem ideal for studying naturalistic forms of threat and safety learning. However, this approach has not yet been exploited.

In this study, we implemented a novel mouse model that involved spatially-distributed thermal reinforcers to examine threat and safety learning. Optogenetic inhibition was performed to evaluate the individual roles of the PL and IL regions during normal conditions. Then, stress treatments were implemented to evaluate changes in PL and IL function during stress-induced disease-like states.

## Materials and Methods

### Subjects

All procedures were approved by the Institutional Animal Care and Use Committee in compliance with the U.S. Public Health Service’s Policy on Humane Care and Use of Laboratory Animals (PHS policy), the Guide for the Care and Use of Laboratory Animals, and the Society’s Policies on the Use of Animals in Neuroscience Research. A total of 299 C57BL/6J male mice were included in this study. They were acquired through a commercial supplier (Jackson Laboratory) and were approximately 8-weeks of age on arrival. Mice were housed in polycarbonate mouse cages (4 per cage) in a vivarium with controlled temperature and pressure, and a 12-hr light/dark cycle with lights on at 7:00 A.M. Food and water were available *ad libitum*, as long as mice were not undergoing behavioral testing. Mice were allowed to acclimate to the vivarium and group housing conditions for at least two weeks prior to any procedure. Then, mice underwent a 10-min handling session, followed in subsequent days by either behavioral testing, stress exposure, or surgical procedures.

### Stereotaxic Surgery

Surgical procedures were performed for bilateral delivery of viral vectors encoding for light-sensitive proteins and chronic implantation of optical fibers for optogenetic manipulations. All surgical procedures were performed under aseptic conditions using stereotaxic frames (KOPF), isoflurane anesthesia systems (Harvard Apparatus), mouse temperature controllers, and stereoscopes for magnification. Prior to skin incision, pre-emptive analgesia was achieved with local injections of lidocaine and bupivacaine (7-8 mg/kg, each), followed by a subcutaneous injection of sustained-release meloxicam (Melox-SR, 4 mg/kg) to provide analgesia for at least 72 hr. For viral infusions into PL, the following stereotaxic coordinates were used: +1.70 mm A/P, ±0.35 mm M/L, and −2.40 mm D/V, relative to bregma. Optical fibers were then positioned in the dorsal portions of PL using a 10° M/L angle, at the following coordinates: +1.70 mm A/P, ±0.75 mm M/L, and −1.95 mm D/V, relative to bregma. For viral infusions into IL, the following stereotaxic coordinates were used: +1.73 mm A/P, ±0.35 mm M/L, and −2.95 mm D/V, relative to bregma. Optical fibers were then positioned in the dorsal portions of IL using a 20° M/L angle at the following coordinates: +1.73 mm A/P, ±1.25 mm M/L, and −2.30 mm D/V, relative to bregma. Optical implants were secured to the skull using a biocompatible adhesive containing methacrylate resin and 4-META monomer (C&B Metabond; Parkell), and self-cure orthodontic black resin acrylic (Ortho-Jet; Lang Dental). Incisions were then sutured, and mice were allowed to recover from surgery for several hours. During this initial recovery period, mice were placed in clean cages with temperature-regulated warming pads and free access to water and soft food options (e.g., nutritional gel cups and wet mouse chow). Mice were then transferred to the animal vivarium for full recovery. Post-operative care was provided by veterinary personnel and experimenters.

### Optogenetic-Mediated Inhibition

After surgical procedures, a period of 8-12 weeks was allowed for efficient expression of viral vectors. After this period, mice then underwent stress exposure and/or behavioral testing. Viral aliquots were obtained from commercial suppliers (Addgene and UNC Vector Core). Viral vectors were infused using a microsyringe pump system (UMP3T-2; Nanofil with 33-Ga needles; WPI). A volume of 400 nL of the viral-containing medium was infused in each hemisphere at the target sites, at a rate of 100 nL/min. The infusion needles were kept at the target site for an additional ten minutes to allow proper diffusion of viruses. Needles were then slowly withdrawn to prevent back pressure and leakage outside the target sites. For optogenetic-mediated neuronal silencing (“photoinhibition”), several cohorts of mice were transduced in the target areas with serotype-5 adeno-associated viral vectors coding for the outward proton-pump *Halorubrum sodomense* archaerhodopsin, fused to enhanced yellow fluorescent protein, under the calcium/calmodulin-dependent protein kinase-II alpha promoter (AAV_5_-CaMKllα-ArchT3.0-eYFP). For comparisons purposes, other control cohorts were transduced with viral vectors that only encoded for eYFP (AAV_5_-CaMKIIα-eYFP). The optical fiber implants were constructed in-house and consisted of Ø300-µm multimode cores (NA = 0.39; Thorlabs) attached to stainless steel ferrules (Ø1.25-mm OD, 330-µm ID bore, Precision Fiber Products). The ferrule ends were polished until achieving a light transmission efficiency >85%, using a fiber connector micropolisher (SpecPro, Krell Tech). Prior to behavioral testing, mice were acclimated to being connected to fiberoptic patch cords (branching fibers, Ø200-µm cores, NA = 0.22, Doric Lenses), which were attached to optical rotaries (FRJ 1×1 FC/PC, Doric Lenses), which in turn were connected to red-shifted DPSS lasers (200 mW max, 589-nm, OptoEngine). Laser output was moderated by mechanical laser beam shutters (100-Hz SR475 Shutter Heads, SRS), which were controlled with TTL pulses using a hardware/software optogenetic interface (ANY-maze). During behavioral experiments, photoinhibition was performed using constant delivery of laser light. Laser power was set to <5 mW at the tip of the patch cords.

### Social Isolation Stress

Social isolation was used to examine the impact of stress on threat and safety learning during the thermal tasks. While the control groups always remained in group-housing conditions, the stress groups underwent individual housing conditions for twelve days. After this isolation period, mice were returned to group-housing conditions for a period of two weeks prior to testing in the thermal threat and safety tasks. This two-week period was provided to allow animals to re-establish social ranks and interactions with their former cagemates. All groups had free access to food and water, and were maintained in the same room and holding rack in the animal vivarium. Body weights were measured every other day to evaluate changes in weight gain by the stress treatment. No optogenetic manipulations were performed during the social isolation stress period.

### Thermal Threat Task

During this spatial-based threat learning task, mice were allowed to explore a square-shaped acrylic box (30 × 30 × 30 cm) in which distinct parts of the floor had different temperatures. While one of the quadrants had a noxious cold temperature (−5°C, “threat zone”; achieved by placing a mixture of regular and dry ice underneath the acrylic floor), the other three quadrants were maintained at a comfortable temperature (30°C; achieved by placing warming mats underneath the acrylic floor). The stability of these temperatures was frequently monitored using a temperature-sensing infrared camera (FLIR C5) and a digital thermometer (LaserPro LP300). To facilitate spatial-based learning, visual cues were added to the walls of the box to differentiate the threat zone (with plus symbols) from the other zones (vertical bars). The training session lasted for ten minutes, which was sufficient to produce optimal behavioral performance, assessed as the time that mice spent avoiding the cold threat zone. A long-term memory test session lasting for three minutes, in the absence of the cold zone, was conducted twenty-four hours after training. During the test session, avoidance of the quadrant that was previously paired with the cold temperature was indicative of good threat recall. Optogenetic manipulations were performed during the ten minutes of the training session. No laser manipulations were performed during the recall test.

### Thermal Safety Task

During this spatial-based safety learning task, mice were allowed to explore a similar square-shaped acrylic box (30 × 30 × 30 cm) in which one quadrant had a pleasant warm temperature (30°C, “safety zone”), while the other quadrants had an uncomfortable cold temperature (5°C). The cold zones in this task were kept above freezing levels (i.e., >0°C) to reduce physical pain, reduce innate reflexes produced by extreme cold temperatures, and persuade animals to explore the entire arena more often. Yet, temperatures <12°C are still significantly noxious and elicit pain and avoidance in mice (Bautista et al., 2007; Lewis and Griffith, 2022). Visual cues on the walls of the box in this version of the task differentiated the safety zone (with plus symbols) from the other zones (vertical bars). The training session lasted for ten minutes, which was sufficient to produce optimal behavioral performance, assessed as the time that mice spent seeking within the warm safety zone. A long-term memory test session lasting for three minutes, in the absence of the warm zone, was conducted twenty-four hours after training. During the test session, seeking behavior within the same quadrant that was previously paired with the warm temperature was indicative of good safety recall. Optogenetic manipulations were performed during the ten minutes of the training session. No laser manipulations were performed during the recall test.

### Elevated-Plus Maze

The elevated-plus maze assay was used to evaluate anxiety-like behavior. The plus maze apparatus consisted of two open arms (30 × 5 cm) and two close arms (30 × 5 × 15 cm) that intersected at a central zone (5 × 5 cm) and were elevated from a tabletop (40 cm). This apparatus was acquired through a commercial source (San Diego Instruments) and was made out of a beige-colored ABS plastic material for enhanced contrast with mice. This assay lasted for a total of nine minutes and was conducted in a room with moderate lighting levels. For optogenetic experiments, the session was divided into three 3-min epochs with alternating laser status (Off-On-Off). Video capturing was performed to track the position of animals within the apparatus using automated software (ANY-maze). The total time that mice spent in the open arms versus the closed arms was quantified during the different laser epochs. Mice typically avoid the open arms, which are more anxiogenic (Walf and Frye, 2007).

### Open-Field Test

The open-field test assay was used to examine anxiety-like behavior and general locomotion (Seibenhener and Wooten, 2015). Mice were allowed to freely explore a square-shaped arena (40 × 40 × 40 cm) for a total of nine minutes. To examine anxiety, the arena was virtually divided into a center zone (25 × 25 cm) and an outer periphery zone. During optogenetic manipulations, the session was divided into three 3-min epochs with alternating laser status (Off-On-Off). The time that mice spent in the center zone was evaluated as an indicator of anxiety-like behavior (i.e., the center zone is anxiogenic whereas the periphery is safer). To examine locomotor activity, quantifications were made for the speed of movement and total distance traveled by mice during the different epochs.

### Real-Time Place Preference/Aversion

For the real-time place preference/aversion assay (Bimpisidis et al., 2020), mice were allowed to freely explore a rectangular acrylic box (50 × 30 × 22 cm) that was virtually divided into two halves using software (ANY-maze). Mouse entry into one half of the arena resulted in closed-loop photoinhibition (i.e., Laser-On), whereas mouse entry into the other half of the arena resulted in the cessation of photoinhibition (i.e., Laser-Off). Two daily sessions lasting thirty minutes each were conducted for each mouse. The side of the box paired with photoinhibition was counterbalanced between the two days. The total times that mice spent on each side of the box were measured using automated tracking software (ANY-maze). Those times were then added up across days, and difference scores between the Laser-On and Laser-Off sides were calculated for each mouse.

### Software & Statistical Analysis

ANY-maze software was used to control lasers, capture videos, and animal tracking for automated quantifications of behavior. Quantifications included the following: (a) Time that mice spent in distinct zones, (b) Number of entries into distinct zones, (c) Total distance traveled, (d) Average speed of locomotion, and (e) Immobilization/freezing behavior (Blanchard et al., 1986). All data was then processed through GraphPad Prism to generate graphs and perform statistical analyses. Normality was verified using the Kolmogorov-Smirnov test, and results were plotted as mean ± standard error of the mean, otherwise indicated. Statistical differences were evaluated using either one-way ANOVA or two-way repeated measures ANOVA, followed by Bonferroni post-hoc tests to confirm group differences. Multiplicity adjusted *P*-values were considered to account for multiple comparisons. Significance thresholds were set to \**P* < 0.05, \*\**P* < 0.01, \*\*\**P* < 0.001, and \*\*\*\**P* < 0.0001.

### Histology

Mice were deeply anesthetized with isoflurane and transcardially perfused with ice-cold phosphate-buffered saline (1X-PBS) and paraformaldehyde (4%-PFA). Brains were collected, fixed for 24 hours in 4%-PFA, and then were equilibrated for 48 hours in a 4%-PFA/30%-Sucrose solution. Coronal sections were then cut at 40 µm using a sliding microtome (HM430, Thermo Scientific). Brain sections containing the mPFC regions of interest (PL and IL) were then mounted on superfrost microscope slides and immersed in a fluorescence-compatible mounting media containing DAPI (Fluoromount-G, Southern Biotech). Images were then acquired using a fluorescence microscope with automated capturing and stitching capabilities (BX43, Cell Sens, Olympus). Viral vector expression and the location of optical fiber tips were then reconstructed in coronal drawings of the mPFC, based on a mouse brain atlas (Paxinos and Franklin, 2004). Only animals in which the PL and IL regions were targeted individually in both hemispheres were considered for analysis. The anatomical boundaries between PL and IL were determined based on anatomical features and transitions in the shape of cortical layers. Due to either viral leakage or misplacement of optical fibers, 36 mice were excluded from the study.

### Availability of Materials and Data

Data will be made available for scientific use upon request. Viral vector sequences are available through commercial suppliers (e.g., Addgene and UNC Vector Core).

## Results

### Noxious cold temperatures were effective for spatial-based threat learning

Our first goal was to examine whether cold temperatures could produce spatial-based threat memory. For this, we designed a square-shaped arena in which a particular quadrant had an extremely cold temperature (−5°C, “threat zone”), while the other quadrants were kept at a comfortable temperature (30°C) (Fig. 1A, left panel). To facilitate spatial-based learning, visual cues were added to the walls of the apparatus to distinguish the threat zone (with plus signs) from the other zones (with vertical columns). The typical response exhibited by mice was avoidance of the quadrant containing the thermal threat (Fig. 1A, right panel).

**Figure 1.**
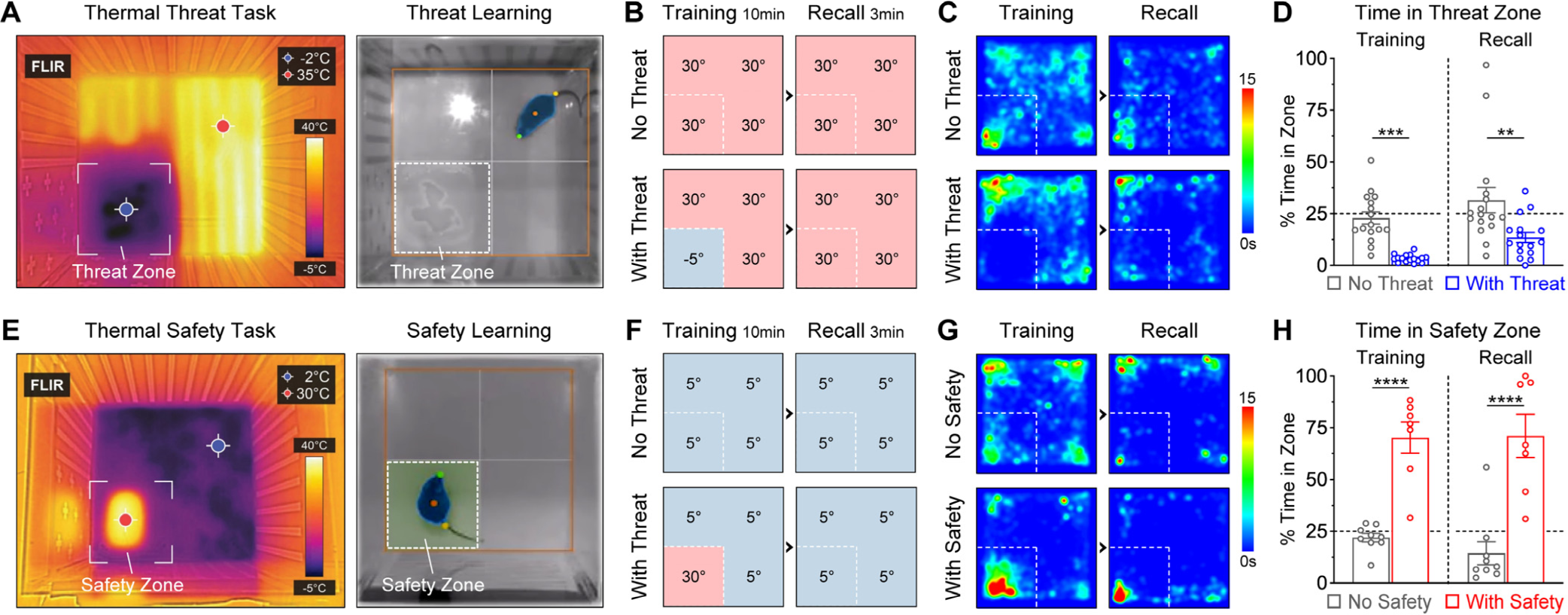
Threat and safety learning tasks using noxious and pleasant temperatures. ***A,*** Thermal threat task. The infrared picture on the left shows merged features in the visible light spectrum. The threat zone was differentiated from the other zones using visual cues on the walls (plus symbols). ***B,*** Schematic for validation of the threat task. The control group never experienced the threat zone (“No Threat”, N = 16), whereas the experimental group experienced thermal threat during training (“With Threat”, N = 16). Both groups were tested the next day in the absence of the threat to evaluate long-term memory recall. ***C,*** Group heatmaps during the threat task, based on the time that mice spent in different locations. ***D,*** Group comparisons during the threat task. Mice that experienced the threat zone continued to avoid the correct quadrant, despite the absence of threat. This indicates good recall of threat memory. ***E,*** Thermal safety task. The safety zone was differentiated from the other zones using visual cues on the walls (plus symbols). ***F,*** Schematic for validation of the safety task. The control group never experienced the safety zone (“No Safety”, N = 9), whereas the experimental group experienced the safety zone during training (“With Safety”, N = 7). Both groups were tested the next day in the absence of safety zone to evaluate long-term memory recall. ***G,*** Group heatmaps during the safety task, based on the time that mice spent in different locations. ***H,*** Group comparisons during the safety task. Mice that experienced the safety zone during training continued seeking safety within the correct quadrant, despite the absence of safety. This indicates good recall of safety memory. [Data shown as mean ± sem. Superimposed scattered dot plots represent individual mice. Dashed line at 25% represents the level of exploration by chance. \*\**P* < 0.01, \*\*\**P* < 0.001, \*\*\*\**P* < 0.0001]

To validate whether this “thermal threat task” was effective to produce a long-lasting memory, comparisons were made between two groups of mice (Fig. 1B). While a “No Threat” control group was allowed to explore the training box in the absence of thermal threat, a “With Threat” experimental group experienced the thermal threat during a training session that lasted ten minutes (Fig. 1B, left panels). The next day, both groups were allowed to re-experience the training box for three minutes in the absence of thermal threat (Fig. 1B, right panels). This allowed us to evaluate long-term recall of threat memory. As appreciated in the group heatmaps (Fig. 1C) and quantifications of time spent in the quadrant of interest (Fig. 1D), the control group exhibited exploration of all quadrants by chance during both sessions (∼25% time spent in each), whereas the experimental group exhibited avoidance to the quadrant paired with thermal threat (∼3% time spent in the quadrant associated with threat). Two-way ANOVA revealed significant group differences (Fig. 1D; Group, *F*_(1,30)_ = 26.3, *P* < 0.0001; Session, *F*_(1,30)_ = 6.99, *P* = 0.013; Interaction, *F*_(1,30)_ = 0.04, *P* = 0.84), and post-hoc tests confirmed that the experimental group spent significantly less time in the quadrant of interest than controls during the training session (*P* = 0.0006), as well as during the recall session (*P* = 0.0015). Additional measurements revealed that this type of learning is independent of the typical freezing responses observed during cue-shock or context-shock paradigms (Extended Fig. 1A-D). These findings indicate that noxious temperatures are highly effective to produce robust threat learning.

### Pleasant warm temperatures were effective for spatial-based safety learning

Using a similar behavioral strategy, we then evaluated whether pleasant warm temperatures could produce spatial-based safety memory. Here, we paired a quadrant of the arena with a pleasant temperature (30°C, “safety zone”), while the rest of the arena had an uncomfortable lower temperature (5°C) (Fig. 1E, left panel). Notice that the uncomfortable noxious temperature here was kept above zero (5°C) to persuade mice to explore the entire arena. Visual cues on the walls of the apparatus distinguished the safety zone (with plus signs) from all other zones (with vertical columns). After some initial exploration of the entire arena, mice typically exhibited approach and seeking behavior within the quadrant with the pleasant temperature (Fig 1E, right panel).

To validate whether this “thermal safety task” was effective for producing a long-lasting safety memory, two groups of mice were compared (Fig. 1F). A “No Safety” control group was allowed to explore the training box in the absence of safety, whereas a “With Safety” experimental group was allowed to experience the safety zone during a ten-minute training session (Fig. 1F, left panels). The next day, both groups were allowed to re-experience the training box in the absence of safety to evaluate long-term memory recall (Fig. 1F, right panels). It can be clearly appreciated in the group heatmaps (Fig. 1G) and quantifications of time spent in the quadrant of interest (Fig. 1H) that the control group exhibited exploration of all quadrants by chance during both sessions (∼25% time of exploration), whereas the experimental group spent the majority of the time within the quadrant associated with the pleasant temperature during both sessions (∼70% time spent in the safety quadrant). Two-way ANOVA detected highly significant group differences (Fig. 1H; Group, *F*_(1,14)_ = 76.6, *P* < 0.0001; Session, *F*_(1,14)_ = 0.23, *P* = 0.64; Interaction, *F*_(1,14)_ = 0.35, *P* = 0.56; Fig. 1H), and post-hoc tests confirmed that the experimental group spent significantly more time in the quadrant of interest than controls during both sessions (both days, *P* < 0.0001). Additional measurements revealed no group differences in freezing behavior (Extended Fig. 1E-H). These findings indicate that pleasant temperatures in the presence of other noxious temperatures are highly effective to produce robust safety learning.

### Threat learning using thermal reinforcers did not require prefrontal activity

The contributions of PL and IL activity for thermal threat learning were examined using an optogenetic photoinhibition approach to silence CaMKIIα-expressing glutamatergic neurons, which are the most abundant cell type in the mPFC and constitute the large majority of outputs from mPFC (Anastasiades and Carter, 2021). PL and IL were targeted individually in different cohorts of mice (Fig. 2A & 2B). While the control groups only expressed the fluorophore eYFP, the experimental groups expressed the proton pump opsin known as ArchT, which produces robust silencing in response to red-shifted light (El-Gaby et al., 2016; Yizhar et al., 2011). Laser light was delivered into the PL or IL regions through chronically implanted optical fibers (full histological reconstruction is shown in Extended Fig. 2).

**Figure 2.**
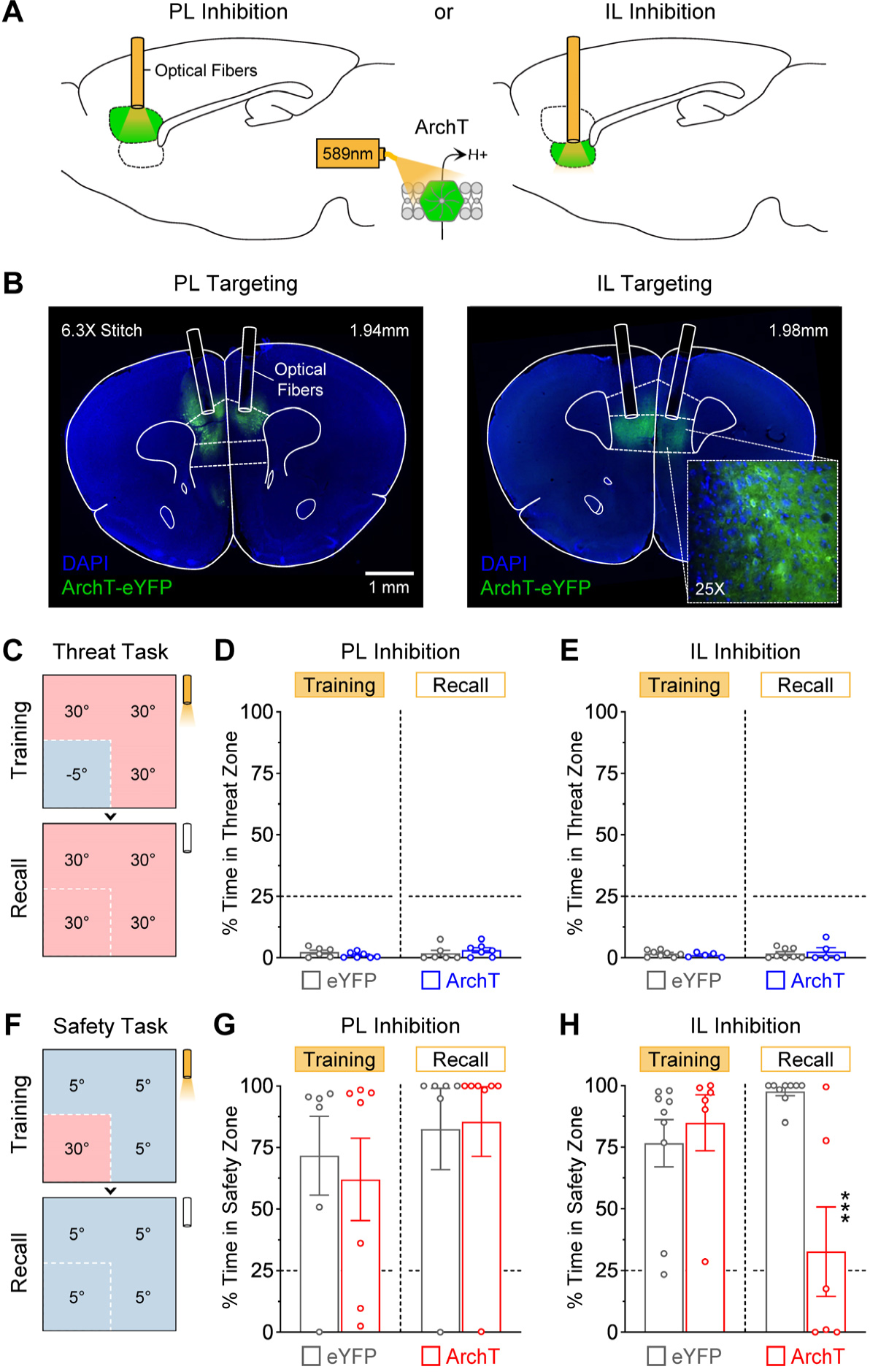
Inhibition of IL (but not PL) impaired safety learning. ***A,*** Strategy to inhibit PL or IL. ***B,*** Representative coronal sections show selective targeting of the PL and IL regions with the viral vectors and optical fibers. ***C,*** Photoinhibition was performed during the thermal threat task. ***D,*** PL inhibition did not affect threat learning (eYFP: N = 6, ArchT: N = 7). ***E,*** IL inhibition did not affect threat learning either (eYFP: N = 8, ArchT: N = 5). ***F,*** In a different cohort of mice, photoinhibition was performed during the thermal safety task. ***G,*** PL inhibition did not affect safety learning (eYFP: N = 6, ArchT: N = 7). ***H,*** IL inhibition did not affect performance during training, but produced a robust impairment in the formation of long-lasting safety memory (eYFP: N = 9, ArchT: N = 6, \*\*\**P* < 0.001).

Photoinhibition was performed throughout the training session of the thermal threat task (Fig. 2C). PL photoinhibition did not affect behavioral performance during training, nor did it affect the formation of a long-term memory during the thermal threat task (Fig. 2D). No significant differences were observed between the PL-eYFP and PL-ArchT groups (Group, *F*_(1,11)_ = 0.0005, *P* = 0.98; Session, *F*_(1,11)_ = 0.54, *P* = 0.48; Interaction, *F*_(1,11)_ = 1.57, *P* = 0.24). Similar to PL, IL photoinhibition neither affected behavioral performance during training, nor affected the formation of a long-term memory during the thermal threat task (Fig. 2E). No significant differences were detected between the IL-eYFP and IL-ArchT groups (Group, *F*_(1,11)_ = 0.005, *P* = 0.51; Session, *F*_(1,11)_ = 0.53, *P* = 0.48; Interaction, *F*_(1,11)_ = 0.47, *P* = 0.51). Thus, thermal threat learning does not require mPFC activity.

### Safety learning using thermal reinforcers required prefrontal activity

In new cohorts of mice, the contributions of PL and IL were examined for thermal safety learning (Fig. 2A & 2B; Extended Fig. 2). As in the previous set of experiments, photoinhibition was performed throughout the training session of the thermal safety task (Fig. 2F). PL photoinhibition neither affected performance during training, nor affected the formation of safety memory (Fig. 2G). Two-way ANOVA confirmed the lack of significant differences between the PL-eYFP and PL-ArchT groups during the safety task (Group, *F*_(1,11)_ = 0.06, *P* = 0.81; Session, *F*_(1,11)_ = 0.89, *P* = 0.37; Interaction, *F*_(1,11)_ = 0.12, *P* = 0.74). On the other hand, while IL photoinhibition did not affect performance during training, it produced a robust impairment in the formation of safety memory (Fig. 2H). In average, the IL-ArchT group exhibited exploratory behavior by chance during the recall session (∼25% time spent in the quadrant previously paired with safety). This is in striking contrast to the IL-eYFP group which exhibited robust recall of the safety memory (∼98% time spent within the quadrant that previously predicted safety). Two-way ANOVA detected significant differences in the group and interaction factors (Group, *F*_(1,13)_ = 5.90, *P* = 0.030; Session, *F*_(1,13)_ = 2.96, *P* = 0.11; Interaction, *F*_(1,13)_ = 16.2, *P* = 0.0014), and post-hoc tests confirmed the lack of group differences during training (*P* > 0.99), but a highly significant difference during recall (*P* = 0.0003). These results indicate that neural activity in IL (but not in PL) promotes safety learning.

### Stress produced major deficits in safety learning but not threat learning during the thermal tasks

It is well-established that uncontrollable stress potentiates threat learning and mimics disease-like states in rodents. Though, it still remains understudied how stress affects naturalistic forms of threat and safety learning. We then examined if learning during our thermal tasks was sensitive to stress. As a stressor, we implemented a social isolation model. While no-stress controls always remained in group-housing conditions (four per cage), mice in the stress groups underwent single-housing conditions for twelve days, followed by group-housing conditions again for an additional two weeks prior to testing in the thermal tasks (Fig. 3A). This period of isolation was more than sufficient to impair body weight gain (Extended Fig. 3), as typically observed for other psychological stressors.

**Figure 3.**
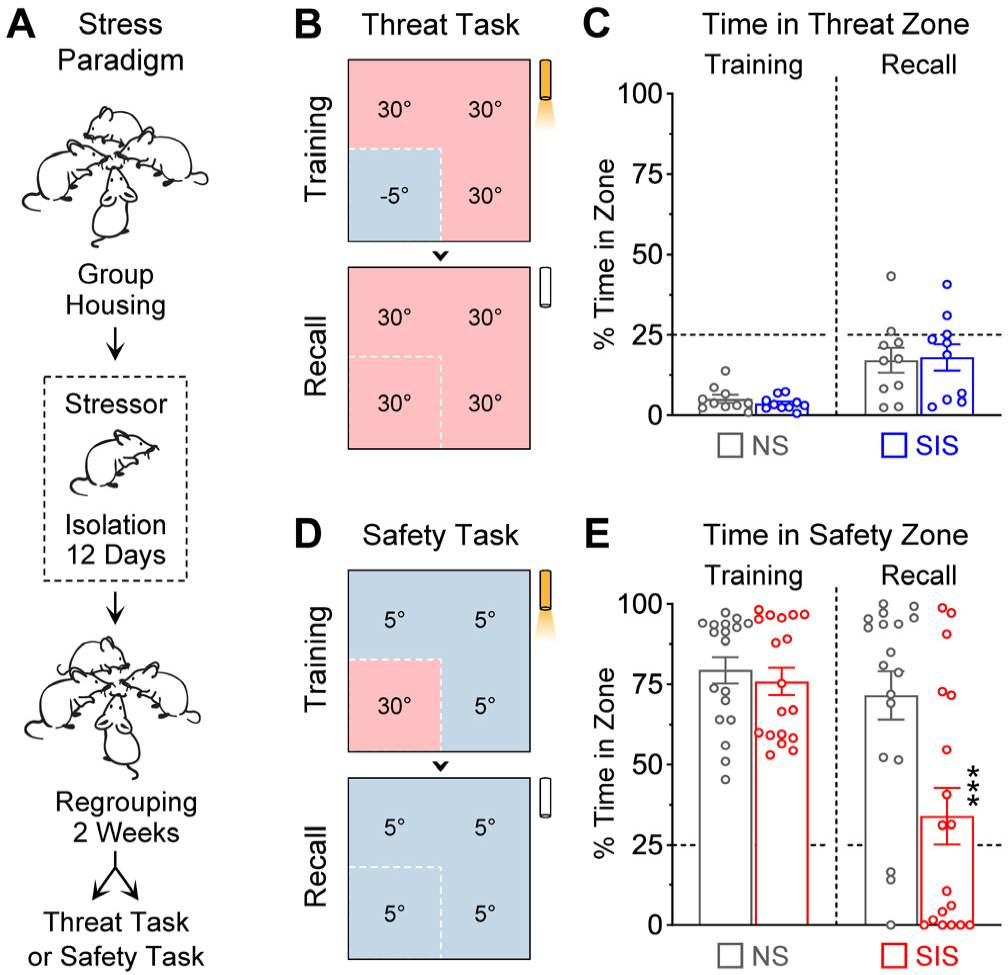
Safety learning was highly susceptible to stress. ***A,*** Schematic of the stress paradigm. While no-stress controls (“NS”) always remained under group-housing conditions, the social isolation stress group (“SIS”) underwent single-housing conditions for twelve days. Two weeks after regrouping, mice underwent thermal threat or thermal safety learning. ***B-C***, In the first cohort, stress did not affect threat learning (NS: N = 10, SIS: N = 10). D-E, In a different cohort, stress-exposed mice showed optimal performance during training, but exhibited a robust impairment in the formation of long-last safety memory (NS: N = 18, SIS: N = 18, \*\*\**P* < 0.001).

Interestingly, during the thermal threat task (Fig. 3B), no effects were produced by stress. Similar to controls, stress-exposed mice exhibited normal behavioral performance and memory formation for the threat quadrant (Fig. 3C). Two-way ANOVA confirmed the lack of significant differences between groups (Group, *F*_(1,18)_ = 0.005, *P* = 0.94; Session, *F*_(1,18)_ = 20.0, *P* = 0.0003; Interaction, *F*_(1,18)_ = 0.15, *P* = 0.70). In contrast, during the thermal safety task (Fig. 3D), strong effects were produced by stress. Compared to controls, the stress-exposed group showed extremely poor safety recall despite exhibiting good performance during the training session (Fig. 3E). Notice that on average, the stress group behaved by chance during the recall session (∼25% time spent in the quadrant previously paired with safety), as if they had never received training. Two-way ANOVA revealed significant differences for the group and interaction factors (Group, *F*_(1,34)_ = 11.2, *P* = 0.002; Session, *F*_(1,34)_ = 13.2, *P* = 0.0009; Interaction, *F*_(1,34)_ = 6.18, *P* = 0.018), and post-hoc tests confirmed the lack of group differences during training (*P* > 0.99), but a highly significant difference during recall (*P* = 0.0002). Thus, while thermal threat learning remained unaffected, thermal safety learning was highly sensitive to stress.

### The stress treatment uncovered bidirectional control of safety learning by the prefrontal regions

We then examined whether additional prefrontal contributions could be uncovered by combining the stress and optogenetic treatments. Several cohorts of mice were prepared for ArchT-mediated inhibition or eYFP-control manipulations in IL or PL (Fig. 4A; histological reconstruction in Extended Fig. 4). Some groups were kept in group-housing conditions (no-stress controls), while other groups underwent social isolation stress (Fig. 4B). Two weeks after the cessation of isolation stress, all groups underwent testing in the thermal safety task (Fig. 4C).

**Figure 4.**
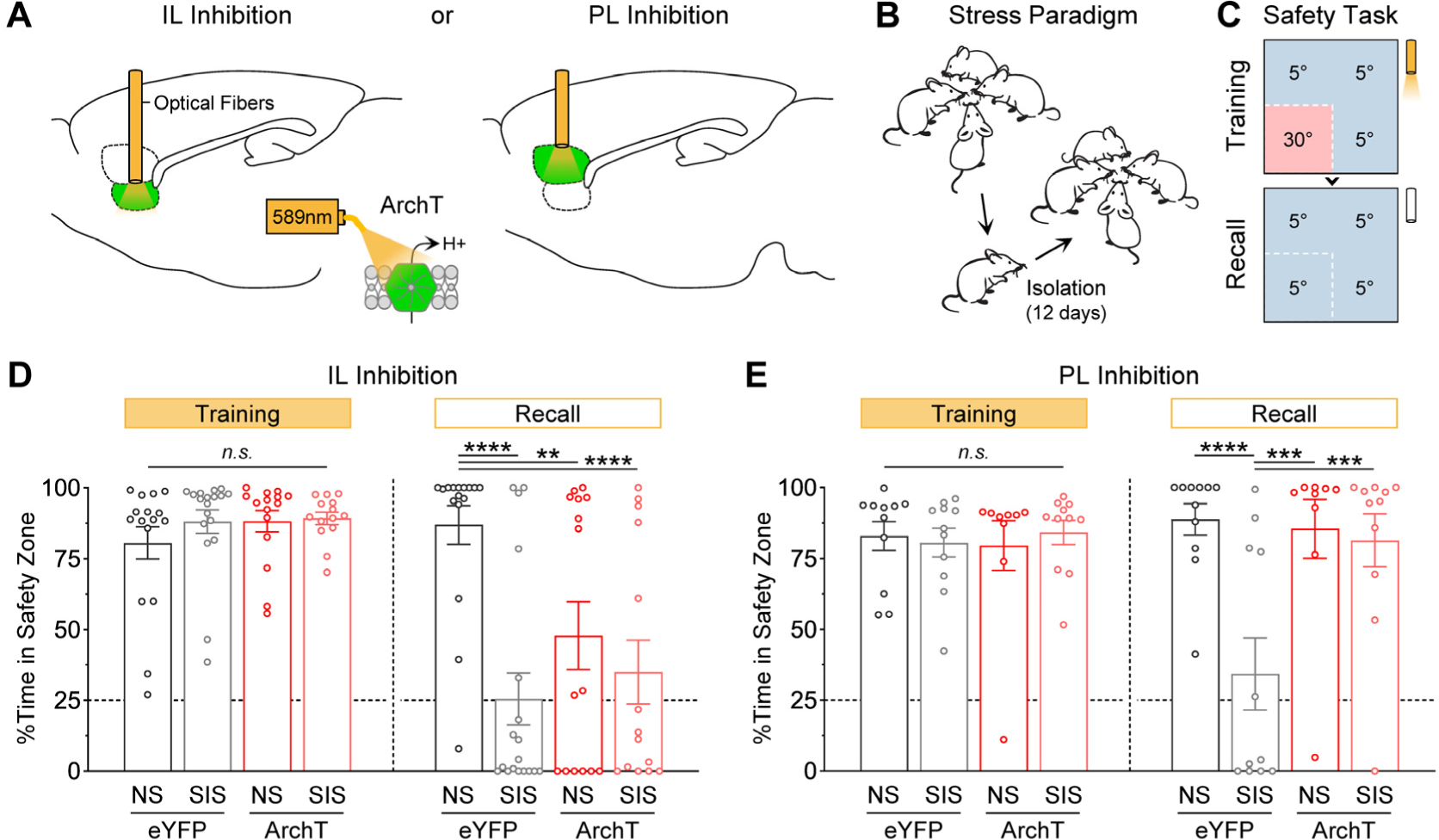
IL inhibition mimicked the effects of stress on safety learning, whereas PL inhibition rescued safety learning in stress-exposed mice. ***A-C***, Schematics for the optogenetic strategy, stress paradigm, and safety learning task. ***D***, IL inhibition produced similar impairment on safety learning to social isolation stress (NS-eYFP: N = 16, SIS-eYFP: N = 18, NSArchT: N = 15, SIS-ArchT: N = 14; \*\**P* < 0.01, \*\*\*\**P* < 0.0001). ***E***, In contrast, PL inhibition fully rescued safety learning in stress-exposed mice (NS-eYFP: N = 11, SIS-eYFP: N = 11, NS-ArchT: N = 9, SIS-ArchT: N = 11; \*\*\**P* < 0.001, \*\*\*\**P* < 0.0001).

IL photoinhibition did not affect behavioral performance during the training session in any group (Fig. 4D). However, during the subsequent long-term test, compared to the double-control group (NS-eYFP, dark-gray), the stress-exposed group that did not undergo IL photoinhibition (SIS-eYFP, light-gray) exhibited poor recall of safety memory. Similarly, the no-stress controls that underwent IL photoinhibition (NS-ArchT, dark-red) also showed poor safety recall. In addition, the double-treatment group (SIS-ArchT, light-red) exhibited poor safety recall. Two-way ANOVA revealed statistical significance in the group and interaction factors (Group, *F*_(3,59)_ = 4.93, *P* = 0.004; Session, *F*_(1,59)_ = 48.9, *P* < 0.0001; Interaction, *F*_(3,59)_ = 8.48, *P* < 0.0001). While post-hoc tests confirmed the lack of group differences during the training session (all comparisons, *P* > 0.99), they revealed highly significant differences during the recall test between the NS-eYFP and SIS-eYFP groups (*P* < 0.0001), NS-eYFP and NS-ArchT groups (*P* = 0.0027), and NS-eYFP and SIS-ArchT groups (*P* < 0.0001). These results suggest that stress produces impairment in safety learning due to significant reductions in IL activity. This idea is consistent with previous reports showing that stress diminishes IL activity (Lee et al., 2011; Park et al., 2021).

PL photoinhibition did not affect behavioral performance during the training session in any group (Fig. 4E). However, during the subsequent long-term test, compared to the double-control group (NS-eYFP, dark-gray), the stress-exposed group that did not undergo PL photoinhibition (SIS-eYFP, light-gray) exhibited poor safety recall. Consistent with the previous experiment not involving stress, the no-stress group that underwent PL photoinhibition (NS-ArchT, dark-red) showed normal safety recall. Finally, to our surprise, the stress-exposed group that underwent PL photoinhibition (SIS-ArchT, light-red) exhibited normal safety recall, indicating that there was a full rescue effect on safety learning in these stress-exposed mice. Two-way ANOVA revealed statistical significances in the group and interaction factors (Group, *F*_(3,38)_ = 6.21, *P* = 0.0015; Session, *F*_(1,38)_ = 2.45, *P* = 0.31; Interaction, *F*_(3,38)_ = 4.49, *P* = 0.0086). While post-hoc tests confirmed the lack of group differences during the training session (all comparisons, *P* > 0.99), they revealed highly significant differences during the recall test between the NS-eYFP and SIS-eYFP groups (*P* < 0.0001), SIS-eYFP and NS-ArchT groups (*P* = 0.0002), and SIS-eYFP and SIS-ArchT groups (*P* = 0.0004). These results are highly intriguing because contrary to IL activity which seems to be promoting safety learning, PL activity seems to be suppressing safety learning, especially after stress. In addition, these findings are consistent with previously suggested notions that stress alters IL and PL activity in opposite manners to produce impairment in safety learning (Lee et al., 2011).

### Photoinhibition did not produce unspecific effects on anxiety, locomotion, or place preference

The mPFC anxiety-like behaviors in rodents (Adhikari et al., 2011; Felix-Ortiz et al., 2016). This raises the question of whether the effects observed in safety learning could be attributed to alterations in anxiety. To evaluate this possibility, we conducted additional behavioral assays on the very same groups of mice used during safety learning. During an elevated-plus maze test (Fig. 5A), anxiety levels were evaluated as the percentage of time that mice spent in the open arms of the maze. No significant differences on anxiety were detected across groups with PL photoinhibition (Fig. 5B; Two-way ANOVA: Group, *F*_(3,38)_ = 0.15, *P* = 0.93; Laser Epoch, *F*_(2,76)_ = 7.32, *P* = 0.0021; Interaction, *F*_(6,76)_ = 1.09, *P* = 0.38; Post-hoc tests for the Laser-On epoch, all *P’s* > 0.99). No significant differences on anxiety were detected across groups with IL photoinhibition either (Fig. 5C; Two-way ANOVA: Group, *F*_(3,59)_ = 0.22, *P* = 0.89; Laser Epoch, *F*_(2,118)_ = 5.94, *P* = 0.0048; Interaction, *F*_(6,118)_ = 0.39, *P* = 0.47; Post-hoc tests for the Laser-On epoch, all *P’s* > 0.75). During an open-field test (Fig. 5D), anxiety levels were evaluated as the percentage of time that mice spent in the center zone of the arena. No significant differences on anxiety were detected across groups with PL photoinhibition (Fig. 5E; Two-way ANOVA: Group, *F*_(3,38)_ = 0.70, *P* = 0.56; Laser Epoch, *F*_(2,76)_ = 3.45, *P* = 0.064; Interaction, *F*_(6,76)_ = 0.71, *P* = 0.65; Post-hoc tests for the Laser-On epoch, all *P’s* > 0.55). No significant differences on anxiety were detected across groups with IL photoinhibition either (Fig. 5F; Two-way ANOVA: Group, *F*_(3,59)_ = 0.79, *P* = 0.51; Laser Epoch, *F*_(2,118)_ = 6.08, *P* = 0.0044; Interaction, *F*_(6,118)_ = 1.71, *P* = 0.13; Post-hoc tests for the Laser-On epoch, all *P’s* > 0.65). Thus, the photoinhibition treatments in this study were insufficient to affect anxiety, and thereby this possibility was ruled out to explain the effects on safety learning.

**Figure 5.**
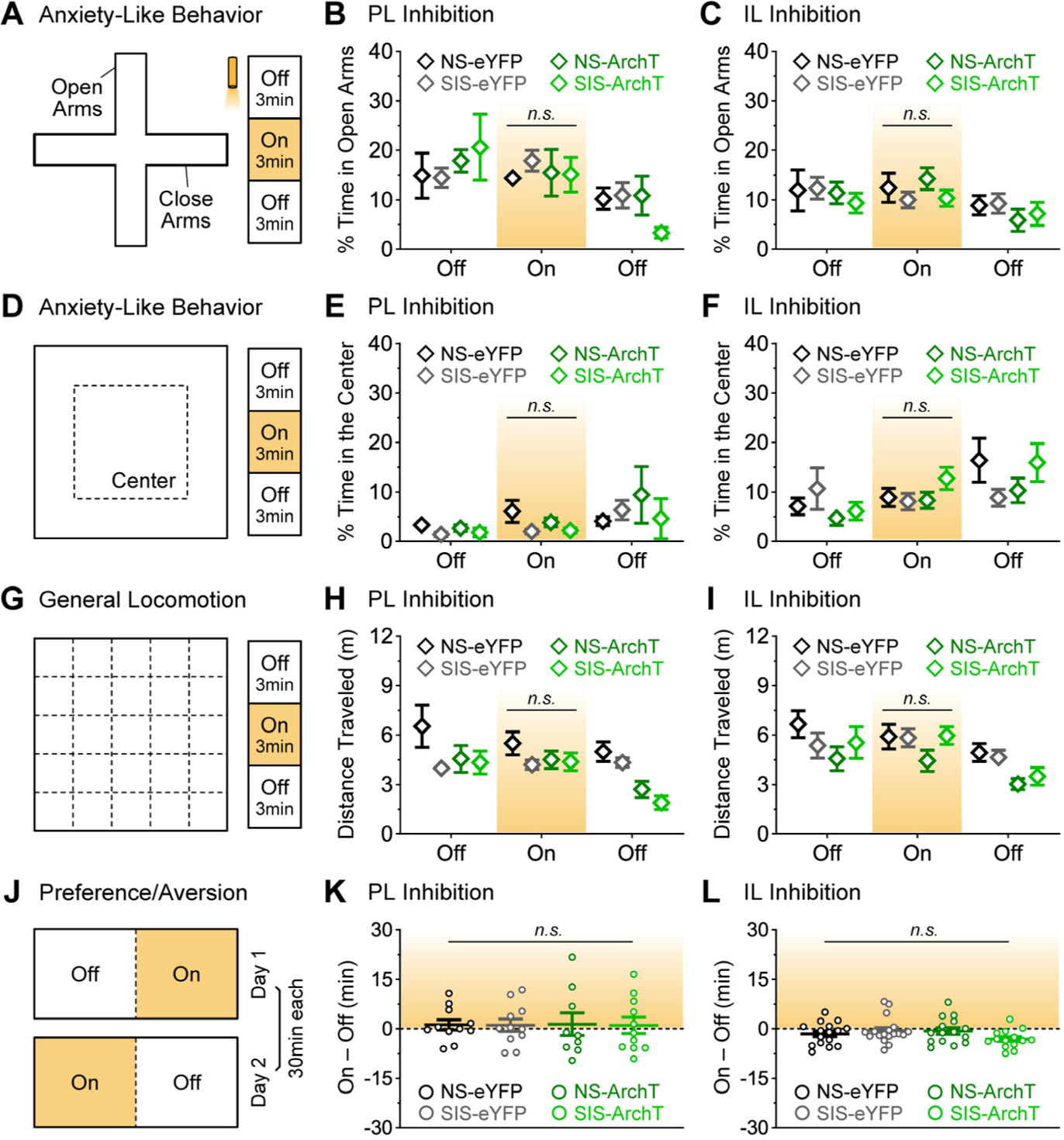
PL and IL photoinhibition did not produce changes in anxiety, locomotion, preference, or aversion. ***A-C***, Anxiety-like behavior in the elevated-plus maze. ***D-F***, Anxiety-like behavior in the open-field test. ***G-I***, General locomotor activity in the open-field test. ***J-L***, Real-time place preference or aversion test. [Number of mice in the PL groups: NS-eYFP: N = 11, SIS-eYFP: N = 11, NS-ArchT: N = 9, SIS-ArchT: N = 11. Number of mice in the IL groups: NS-eYFP: N = 16, SIS-eYFP: N = 18, NS-ArchT: N = 15, SIS-ArchT: N = 14]

Another potential confounding factor could be non-specific effects in general locomotor activity. This possibility was examined during the open-field test in which locomotion was measured as the cumulative distance that mice traveled in the arena (Fig. 5G). No significant differences were detected on distance traveled across groups with PL photoinhibition (Fig. 5H; Two-way ANOVA: Group, *F*_(3,38)_ = 4.17, *P* = 0.012; Laser Epoch, *F*_(2,76)_ = 7.62, *P* = 0.0033; Interaction, *F*_(6,76)_ = 1.92, *P* = 0.089; Post-hoc tests for the Laser-On epoch, all *P’s* > 0.63). No significant differences on distance traveled were detected across groups with IL photoinhibition either (Fig. 5I; Two-way ANOVA: Group, *F*_(3,59)_ = 1.98, *P* = 0.13; Laser Epoch, *F*_(2,118)_ = 14.9, *P* < 0.0001; Interaction, *F*_(6,118)_ = 0.89, *P* = 0.50; Post-hoc tests for the Laser-On epoch, all *P’s* > 0.47). Thus, locomotor effects are ruled out to explain effects on safety learning.

Finally, optogenetic treatments on their own could elicit place preference or aversion, depending on whether they are capable of producing rewarding or noxious effects, respectively. To evaluate this possibility, a real-time place preference or aversion assay was conducted in which the photoinhibition treatments were performed in a closed-loop manner when mice explored one half of a rectangular-shaped box, but not when exploring the other half (Fig. 5J). Place preference or aversion were not observed with PL photoinhibition (Fig. 5K; One-way ANOVA: *F*_(3,38)_ = 0.005, *P* > 0.99). Such effects were not observed with IL photoinhibition either (Fig. 5L; One-way ANOVA: *F*_(3,59)_ = 1.68, *P* = 0.18). Therefore, this possibility can also be ruled out to explain the effects on safety learning.

## Discussion

Safety learning is an essential function for adaptive behavior and mental health. While safety learning is affected in stress-related disorders, the mechanisms normally controlling this function and how they become affected during stress conditions still remain not well understood. New insights were obtained in this study while implementing a semi-naturalistic behavioral paradigm approach using spatially-distributed thermal reinforcers. While focusing on discrete divisions of the mPFC, we found that neural activity in the IL region promoted safety learning during normal conditions, whereas neural activity in the PL region suppressed safety learning after stress. Furthermore, while IL inhibition mimicked the detrimental effects of stress on safety learning, PL inhibition fully restored safety learning in stress-exposed mice. Collectively, these findings highlight the importance of these mPFC regions for bidirectional control of safety learning during normal and stress-induced disease-like states.

### Advantages of studying threat and safety learning using thermal reinforcers

Despite their simplicity, the learning tasks implemented here provide new opportunities to examine the neural substrates of threat and safety learning while mice experience semi-naturalistic conditions. Furthermore, in these learning tasks most mice learned threat and safety with relatively minimal training (within ten minutes), and optimally recalled long-term memory during subsequent test sessions in the absence of the thermal reinforcers. These findings indicate that strong spatial-based memories for threat and safety can be rapidly established with semi-naturalistic settings involving thermal reinforcers. Robust fast learning in these tasks is indeed a tremendous advantage, compared to other types of paradigms that are typically implemented for studying threat and safety. For example, during associative tasks, substantial training is required for optimal dissociation of the cues that differentially predict shock (threat) versus no shock (safety) (Meyer and Bucci, 2014; Rescorla, 1969; Sangha et al., 2013). Another advantage of the thermal tasks is that threat and safety learning produced distinctive adaptive behaviors. While threat learning was represented by avoidance behavior, safety learning was represented by seeking behavior. In contrast, during cue discrimination tasks, while threat learning is represented by conditioned freezing responses to the shock-predicting cues, safety learning is rather inferred from the lack or reduction of freezing responses during the cues that predict the omission of shock (Meyer and Bucci, 2014; Rescorla, 1969; Sangha et al., 2013). Thus, the present tasks provide new chances for studying threat and safety from a more naturalistic perspective in fast and easily reproducible ways.

### Potential disadvantages of the thermal threat and safety tasks

A potential disadvantage of the thermal learning tasks, as implemented in this study, is that they do not include a trial-based structure. This could introduce some challenges when evaluating the neural correlates mediating these types of learning, for example while using in vivo electrophysiology or calcium imaging techniques, due to potential low sampling (i.e., less than ten minutes of training) or due to a reduced number of instances in which animals perform the behaviors of interest (e.g., in average our mice exhibited exploration of either the threat zone or the safety zone less than ten times during training). To overcome these challenges, future studies could either extend the training session, perform multiple daily training sessions, or implement a trial-based design in which mice are removed and reintroduced into the test arena multiple times during a single day, as achieved in other spatial learning tasks (Morris, 1984; Vorhees and Williams, 2006). Another possible strategy is to occasionally drop food pellets in different parts of the arena to persuade animals to explore the threat zone more often or to momentarily leave the safety zone to gather food. Such scenarios may, however, create even more complex situations involving multiple naturalistic motivational conflicts. This could actually allow new avenues of research for multi-dimensional assessment of adaptive behavior from a naturalistic perspective (Bravo-Rivera and Sotres-Bayon, 2020; Fernandez-Leon et al., 2021; Wang et al., 2021).

### Lack of PL and IL involvement during thermal threat learning

Despite our initial predictions that at least the PL region may play an important role, inhibition of neither the PL nor IL region affected performance or learning during the thermal threat task. Our predictions were based on previous evidence that neuronal ensembles in PL exhibit strong representations for the acquisition and expression of threat-related behaviors, including freezing and avoidance (Bravo-Rivera et al., 2015; Burgos-Robles et al., 2017, 2009; Corches et al., 2019; Diehl et al., 2018; Gilmartin and McEchron, 2005; Rozeske et al., 2018). Furthermore, PL inactivation significantly impairs threat-elicited behaviors (Bravo-Rivera et al., 2014; Corcoran and Quirk, 2007; Gilmartin and Helmstetter, 2010; Sierra-Mercado et al., 2011). However, these PL contributions have been mostly reported during tasks in which behavioral responses to environmental predictors of threat emerge as a consequence of recently acquired experiences. In contrast, our threat task involved a naturalistic thermal reinforcer, and behavioral responses were likely driven by innate defensive reflexes associated to pain signals produced by the extremely low temperature within the threat zone (−5°C). Such defensive reflexes are largely driven by hardwired stimulus-response circuits that bypass cortical processing for the rapid triggering of defensive reactions (LeDoux and Daw, 2018). These same reasons could also explain why stress also failed to produce effects during our thermal threat task, whereas stress has been shown to produce strong enhancement of threat learning during cue-shock associative tasks (Conrad et al., 1999; Corley et al., 2012; Maren and Holmes, 2016; Rau and Fanselow, 2009).

### Contributions of IL and PL on safety learning

The mPFC, in particular the IL region, has been implicated in multiple forms of safety learning. For instance, IL activity is needed for the extinction of threat-related responses when environmental cues no longer predict shock punishment (Sierra-Mercado et al., 2011). IL activity also keeps threat responses low during the presentation of safety cues (i.e., stimuli that predict the lack of shock punishment) (Kreutzmann et al., 2020; Sangha et al., 2014). Similarly, IL activity suppresses threat responses when safety cues are presented simultaneously with shock-predicting cues (i.e., conditioned inhibition; Meyer and Bucci, 2014; Sangha et al., 2014). In addition, it has been reported that significant neuronal ensembles in IL signal extinguished cues, safety cues, as well as safe contexts (Corches et al., 2019; Milad and Quirk, 2002; Ng and Sangha, 2022). Collectively, these previous observations suggest that information processing in IL is an essential component for controlling threat responses and promoting adaptive behavior during different forms of safety learning. The present findings expand this view by showing that IL activity is also essential for spatial-based safety learning during situations involving naturalistic thermal threat. In addition, while the impact of stress on safety learning remained largely understudied, we showed that stress pre-exposure (in the form of social isolation) produced robust deficits in safety learning during thermal threat. Furthermore, we showed that selective inhibition of the IL region was sufficient to fully mimic the stress deficits in safety learning. These new observations further highlight the importance of IL activity for promoting safety learning.

In contrast to IL, PL activity seemed to suppress safety learning. This effect became more evident after stress, when selective inhibition of the PL region fully restored safety learning in stress-exposed mice. While we particularly implemented a social isolation stress paradigm, it has been reported that this as well as other stressors (e.g., restraint) are capable of significantly modulating baseline neuronal activity and intrinsic excitability in the PL and IL regions. While PL shows elevated activity, IL shows suppressed activity with stress (Lee et al., 2011; Park et al., 2021). These differential impacts of stress on PL and IL activity seem to produce an unbalanced system that deeply affects safety learning.

### Model of balanced activity between PL and IL, and implications for stress disorders

We propose a model in which balanced activity between the PL and IL regions promotes optimal safety learning (Fig. 6A). However, stress can tilt this balance off (PL > IL), thus producing significant impairment in safety learning (Fig. 6B). Manipulations of this PL-IL balance in healthy controls (e.g., with optogenetic inhibition of IL) lead to an unbalanced system that mimics the impact of stress on safety learning (Fig. 6C). Finally, manipulations capable of restoring the PL-IL balance in stress-exposed subjects is sufficient to rescue safety learning (Fig. 6D).

**Figure 6.**
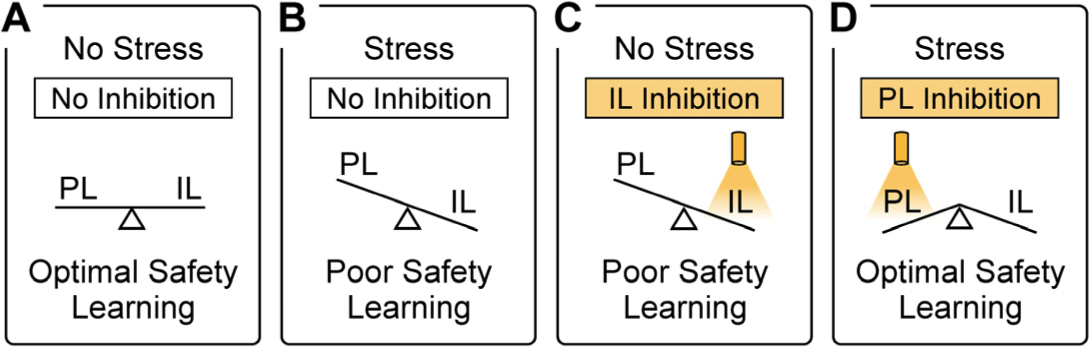
Model of balanced activity between PL and IL for controlling safety learning during health and disease. ***A***, Without stress, balanced activity between PL and IL is suitable for optimal safety learning. ***B***, Consistent with published results (Lee et al., 2011), stress exposure tilts this balance off by increasing PL activity while decreasing IL activity, resulting in poor safety learning. ***C***, IL photoinhibition also tilts off the balance between PL and IL, thus producing poor safety learning in unstressed controls. ***D***, Under stress conditions, PL photoinhibition restores the balance in this prefrontal system, thus rescuing optimal safety learning in stress-exposed mice.

Consistent with our prefrontal balance model and stress-induced alterations, unbalanced prefrontal systems seem to also be characteristic in post-traumatic stress disorder (PTSD). For instance, the dorsal anterior cingulate cortex (dACC), which is a functional homologue to the rodent PL, exhibits enhanced resting metabolic activity and potentiated activation during threat cues in PTSD (Marin et al., 2016; Milad et al., 2007; Shin et al., 2009). In contrast, the ventromedial prefrontal cortex (vmPFC), which is a functional homologue to the rodent IL, exhibits reduced resting activity and diminished activation during extinction of threat responses in PTSD (Marin et al., 2016). While these prefrontal pathologies have been mostly interpreted in terms of dysregulated responses to threat, the observations obtained in the present study encourage further translational research to continue disentangling the mechanisms contributing to threat versus safety learning in humans. If consistent roles are obtained for the dACC and vmPFC for controlling safety learning, the present findings suggest that potential interventions aimed at restoring the balance of activity in these prefrontal systems could be beneficial for improving safety learning, facilitate control over environmental threats, and improve mental well-being in stress disorders.

Finally, while the present study provides novel insights on the contribution of mPFC regions during safety learning, only male mice were considered. It is important to recognize that the prevalence of stress disorders is higher in women than men (McLean et al., 2011). Furthermore, significant sex differences have been documented during other animal models of safety learning (Clark et al., 2019; Krueger and Sangha, 2021; Trask et al., 2020). Thus, future studies could further examine sex differences in safety learning using the paradigms implemented in the present study.

## Acknowledgements

This work was supported by the College of Science at UTSA and a UT-System program for Science and Technology Acquisition and Retention (STARs). Support was also provided by NIH grants R25EB027605, T34GM145507, and T34GM008073. The authors thank Danny Hajali, Gianna Aleman, Daniel Arriaga, and Bianca Moreno for technical assistance. The authors also thank Dr. Jose Rodriguez-Romaguera (UNC) for insightful comments on the manuscript.

## Author Contributions & Conflict of Interest

ACFO and ABR designed the research, analyzed the data, and wrote the paper. ACFO, JMT, CG, HDM, ARR, MBB, SMLP, GM, and ABR performed the research. The authors declare no conflict of interest.

---

**Extended Figure 1.**
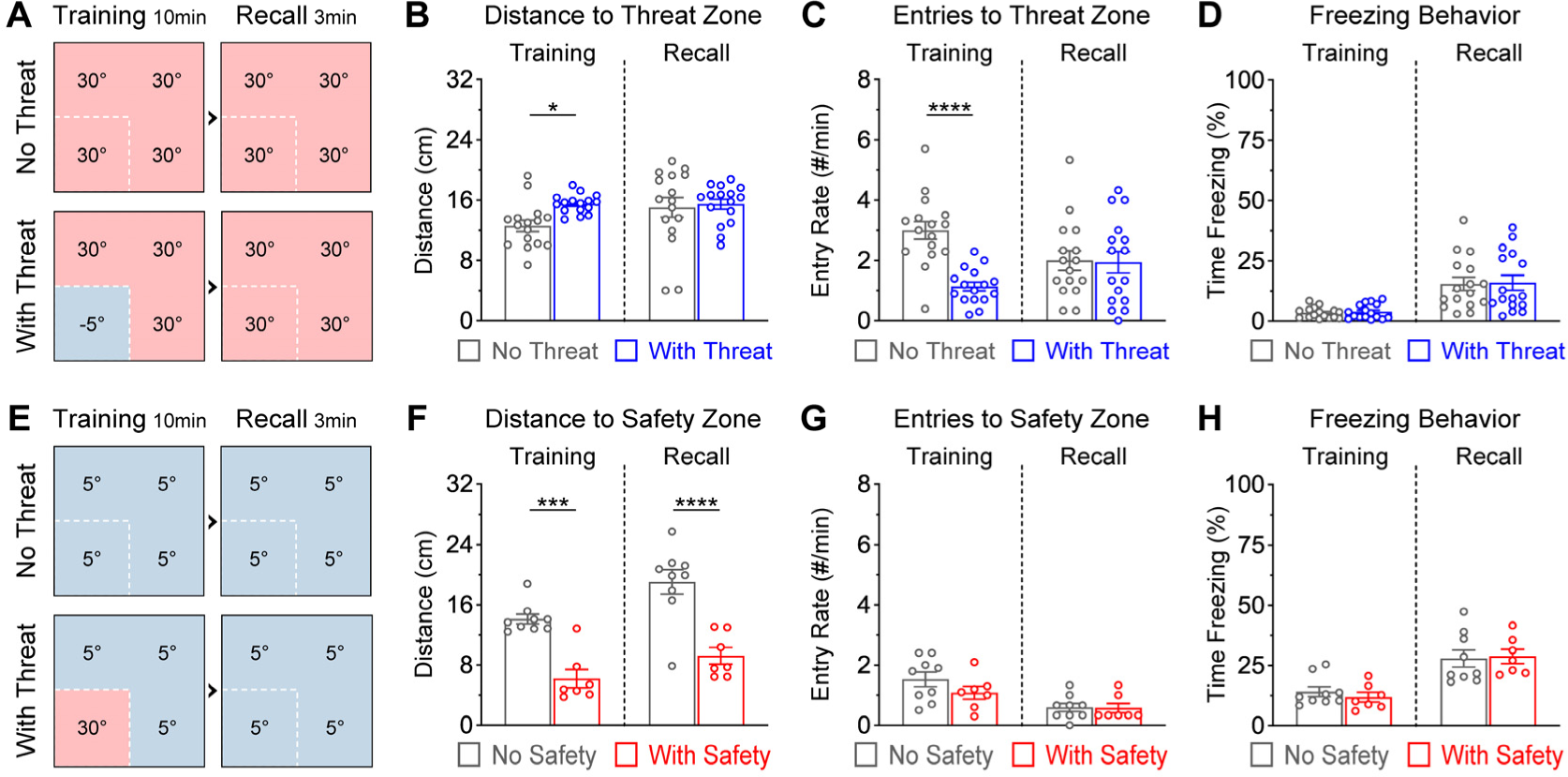
Supporting data analyses for validation of the thermal tasks. ***A,*** Diagram of the threat task. ***B,*** Mice in the experimental group maintained a slightly greater but significant distance between them and the cold threat zone, compared to mice in the control group to the corresponding zone, during the training session (\**P* = 0.034). ***C,*** The experimental group also exhibited a significantly lower number of entries into the cold threat zone during training (\*\*\*\**P* < 0.0001). ***D,*** No group differences were detected on freezing/immobilization levels, which in general remained relatively low during the threat task. ***E,*** Diagram of the safety task. ***F,*** The experimental group maintained a significantly smaller distance between them and the warm safety zone, compared to controls for the corresponding zone, during both sessions (\*\*\**P* = 0.0002, \*\*\*\**P* < 0.0001). ***G,*** No group differences were detected on the number of entries into the warm or corresponding zone. ***H,*** No group differences were detected on freezing/immobilization levels during the safety task either. [The *P*-values reported in this figure represent Bonferroni post-hoc tests after two-way repeated measures ANOVA tests.]

**Extended Figure 2.**
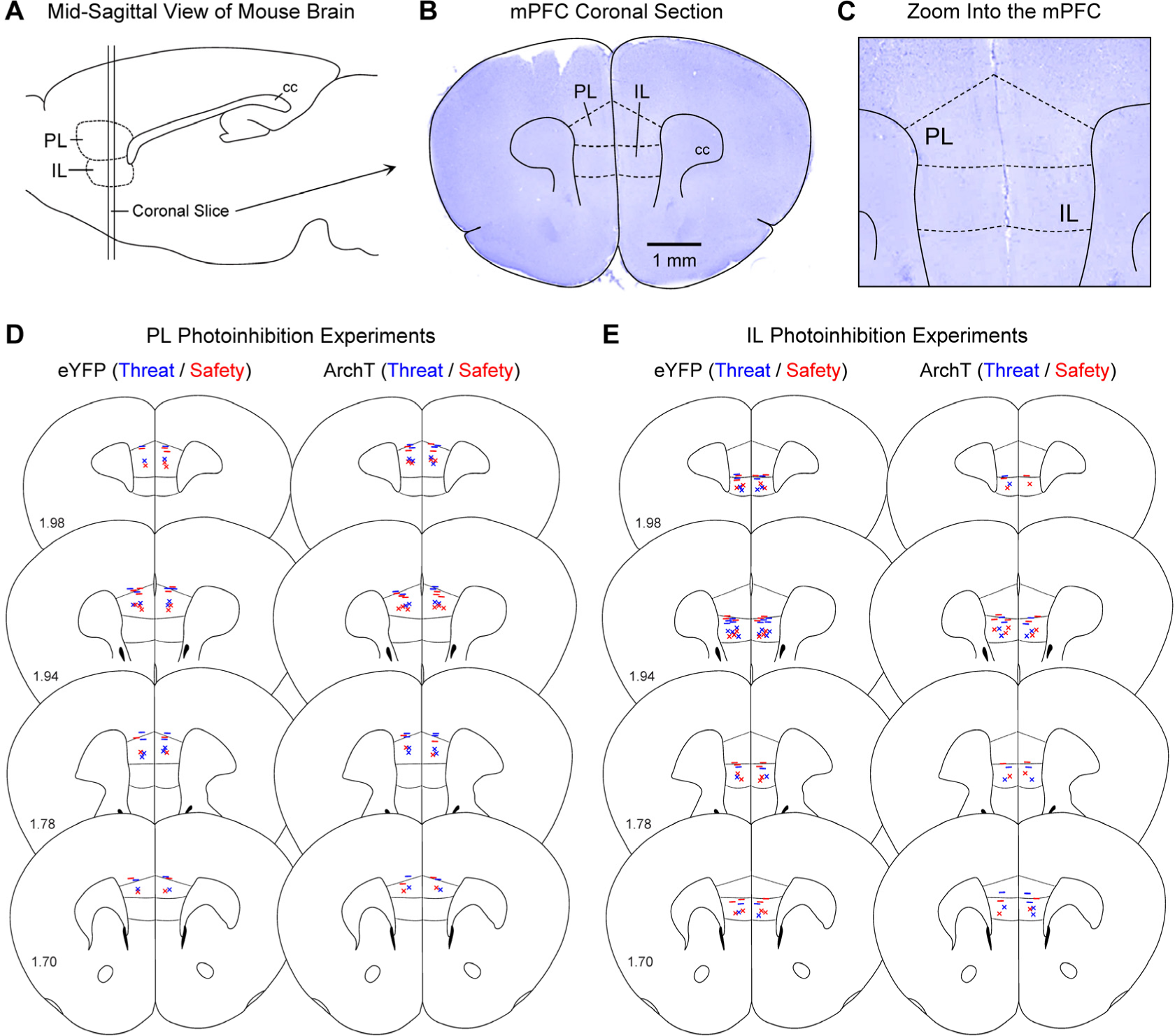
Viral infusion sites and optical fiber placements. ***A,*** Mid-sagittal drawing of the mouse brain illustrating the regions of interest in the mPFC. ***B,*** Coronal photomicrograph of the mPFC. ***C,*** Zoom into the mPFC regions. The shape and transition of cortical layers, as well as other tissue landmarks and cytoarchitectonic features were carefully inspected in the tissue from every mouse to separate the PL and IL regions. ***D-E,*** Summary of the position of optical fiber tips (represented with the “–” symbols) and the centroid of viral vector infusions (represented with the “×” symbols). Numbers within the coronal drawings indicate the anterior-posterior coordinates in millimeters, relative to bregma. [eYFP: control groups, ArchT: neural inhibition groups, cc: corpus callosum]

**Extended Figure 3.**
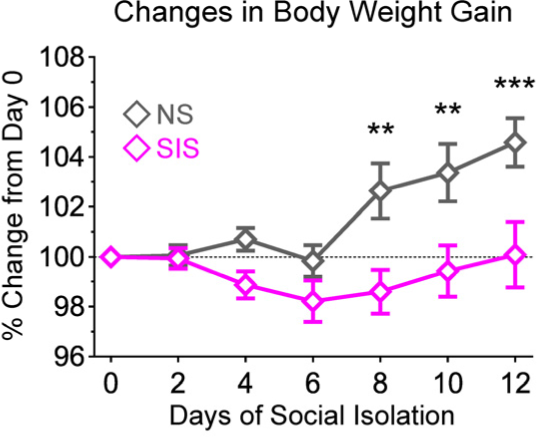
Effects of social isolation on body weight gain. Body weight gain normalized to Day 0 (\*\**P* < 0.01, \*\*\**P* < 0.001).

**Extended Figure 4.**
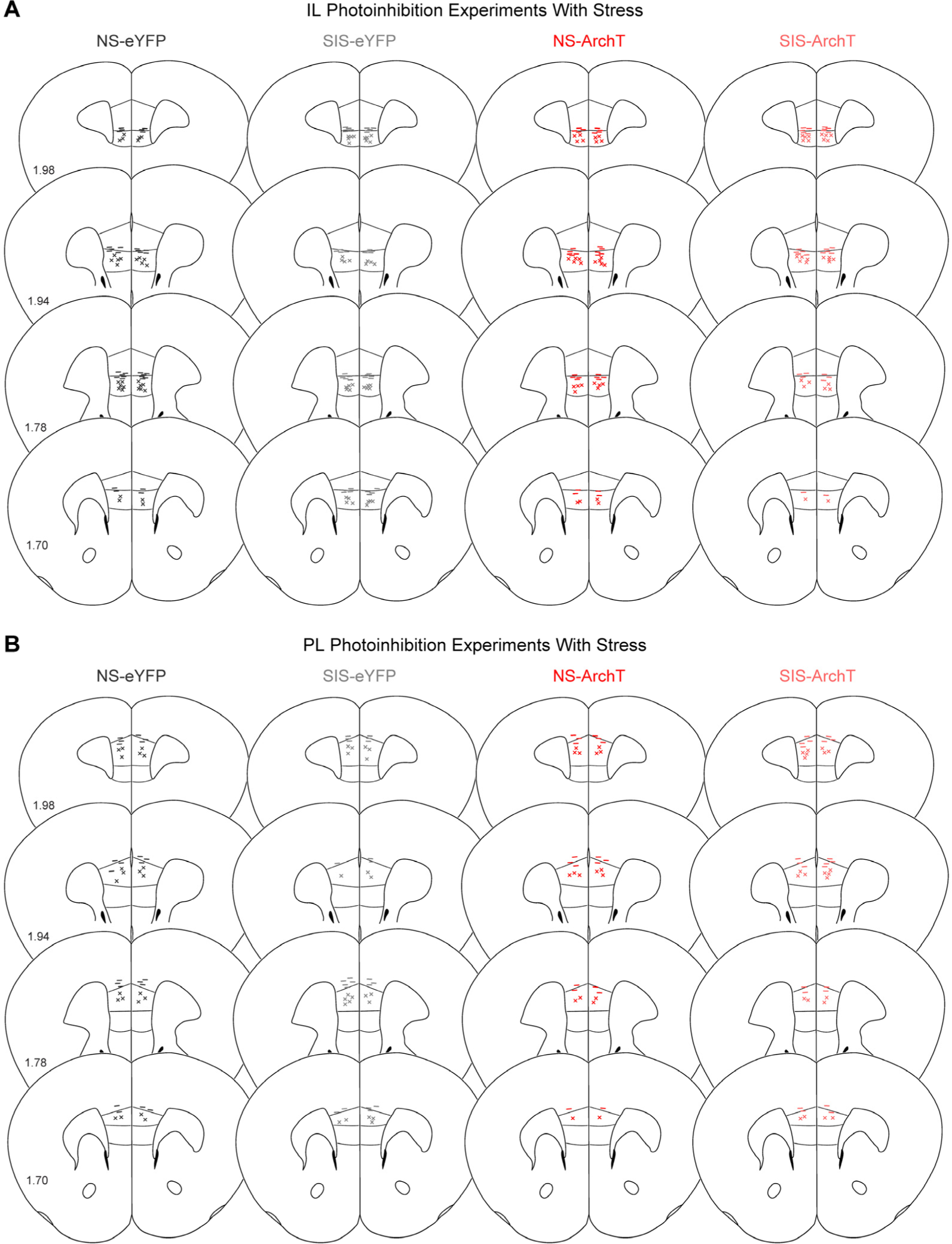
Viral infusion sites and optical fiber placements for the experiments implementing stress. ***A***, IL. ***B***, PL. The “–” symbols represent the location of optical fiber tips and the “×” symbols represent the centroid of viral vector infusion. [NS: no stress, SIS: social isolation stress, eYFP: control fluorophore, ArchT: neural inhibition opsin]

